# optimalTAD: annotation of topologically associating domains based on chromatin marks enrichment

**DOI:** 10.1101/2023.03.06.531254

**Authors:** Dmitrii N. Smirnov, Anna D. Kononkova, Debra Toiber, Mikhail S. Gelfand, Ekaterina E. Khrameeva

## Abstract

In many eukaryotes, chromosomes are organized as strings of spatially segregated Topologically Associating Domains (TADs), characterized by a substantially increased frequency of interactions within them. Boundaries of TADs are highly enriched in histone acetylation chromatin marks and occupied binding sites of architectural proteins, highlighting the functional role of TADs in the regulation of gene expression. While many computational approaches have been developed for TAD identification, it remains challenging because of their nested structure, resulting in weakly overlapping sets of TADs at different scales. Here, we propose a novel algorithm `optimalTAD` for identifying the optimal set of TADs based on epigenetic marks enrichment. Assuming that the most dramatic enrichment corresponds to the best annotation of TAD boundaries, our algorithm optimizes TAD calling parameters by maximizing the difference in chromatin mark levels between TADs and their boundaries. Using this algorithm, we annotated TADs in multiple publicly available fruit fly and mammalian Hi-C datasets and identified a set of epigenetic marks that are best suited for TAD prediction. Through the analysis of diverse organisms and cell types with distinct underlying principles of TAD organization, we have shown that `optimalTAD` is a universal tool suitable for studying TAD structure, functions, and properties unique to specific cell types and organisms. `optimalTAD` is freely available at GitHub: https://github.com/cosmoskaluga/optimalTAD.

**Key Points:**

- We assume that the most dramatic enrichment of epigenetic marks corresponds to the best annotation of TAD boundaries.
- Our algorithm optimizes TAD calling parameters by maximizing the difference in chromatin mark levels between TADs and their boundaries.
- `optimalTAD` is a universal tool that is applicable for studying TAD characteristics in diverse organisms and cell types.
- `optimalTAD` enables the identification of a specific set of epigenetic marks that are most suitable for annotating TADs.

## Background

Chromatin conformation capture methods (3C and Hi-C) have uncovered a fundamental level of chromosome organization: Topologically Associating Domains (TADs) [7, 24]. The average TAD size differs across species: 1000 kb for the human, 880 kb for the mouse, and 140 kb for the fruit fly genome. Besides different sizes, TAD structural properties also vary between species. While mammalian chromatin is partitioned into contiguous topological domains, TADs in *Drosophila* are separated by stretches of decompressed chromatin - interTADs. According to recent Hi-C experiments with sub-kb resolution, inter-TAD regions are composed of small weak TADs (median size 9 kb) enriched for active chromatin [38]. These small active TADs are indistinguishable at lower resolution levels and therefore cannot be resolved in most Hi-C studies traditionally separating *Drosophila* chromatin into TADs and inter-TADs.

TADs represent functional domains of gene expression regulation, defining promoter-enhancer interactions through chromatin loops [8]. TAD boundaries are enriched in active transcription and housekeeping genes [36], which creates a pronounced gradient of histone acetylation and other active chromatin marks at TAD boundaries. In mammals, TAD boundaries are occupied by insulator proteins (Cohesin, CTCF), maintaining spatial segregation of TADs [13]. In *Drosophila*, other architectural proteins (BEAF32, Chromator) are enriched at TAD boundaries [34]. The reported gradient of chromatin marks between TADs and their boundaries offers valuable information for precise annotation of TADs, yet none of the existing TAD-calling techniques takes it into account.

Several TAD-calling algorithms have been introduced, including `Directionality Index` [7], `Armatus` [10], `Arrowhead` [29], `CITD` [5], `HiCseg` [19], `TopDom` [33], `calTADs` [40], `TADtree` [41], `Insulation Score` [27]. Among these methods, the `Armatus` algorithm deserves special recognition because it provides a multiscale approach for TAD calling and can be successfully applied to both mammalian and *Drosophila* genomes [30, 36]. The multiscale nature of `Armatus` allows it to produce different TAD sets depending on the value of the resolution parameter *γ*. TADs in the resulting sets vary substantially in size and number, depending on *γ*. Similarly, the `Insulation Score` approach [6] is based on the insulation profile reflecting the aggregate of interactions within a sliding diamond-shaped genomic window of variable size. Minima of the insulation profile indicates regions of high insulation classified as TAD boundaries. Like in `Armatus`, the size and number of the TADs produced by `Insulation Score` depend on the choice of window size. Such variability might reflect the biological properties of predicted domains but impedes their interpretation because it is not clear which TAD set better matches chromatin functional properties and is therefore suitable for further analysis.

Here, we present a user-friendly and accurate tool called `optimalTAD` to determine the optimal set of TADs based on a combination of Hi-C and epigenetic data for robust annotation of TADs as both structural and functional chromatin units. For a given Hi-C matrix, `optimalTAD` identifies a complete set of multiscale domains via the `Armatus` or `Insulation Score` methods and then evaluates each TAD set using ChIP-seq profiles of histone marks or architectural proteins to obtain the optimal TAD segmentation.

## Results

### The optimalTAD algorithm

The algorithm relies on data collected from both Hi-C and epigenetic experiments, such as ChIP-seq, meDIP-seq, or similar. (Fig. 1). For a given Hi-C contact matrix, `optimalTAD` utilizes the `Armatus` algorithm to derive TAD annotations. Importantly, the `Armatus` average TAD size depends on the value of the resolution parameter *γ*. In our approach, `Armatus` generates multiple TAD sets, with each TAD set corresponding to a different *γ* value ranging from *γ*_min_ *>* 0 to *γ*_max_ with a step *s*. To select the optimal TAD annotation, `optimalTAD` calculates medians *M* ^TAD^ and *M* ^interTAD^ of the scaled epigenetic values within TAD and inter-TAD regions, followed by the maximization of the difference between these medians across the various *γ* values (Fig. 1).

**Fig. 1.**
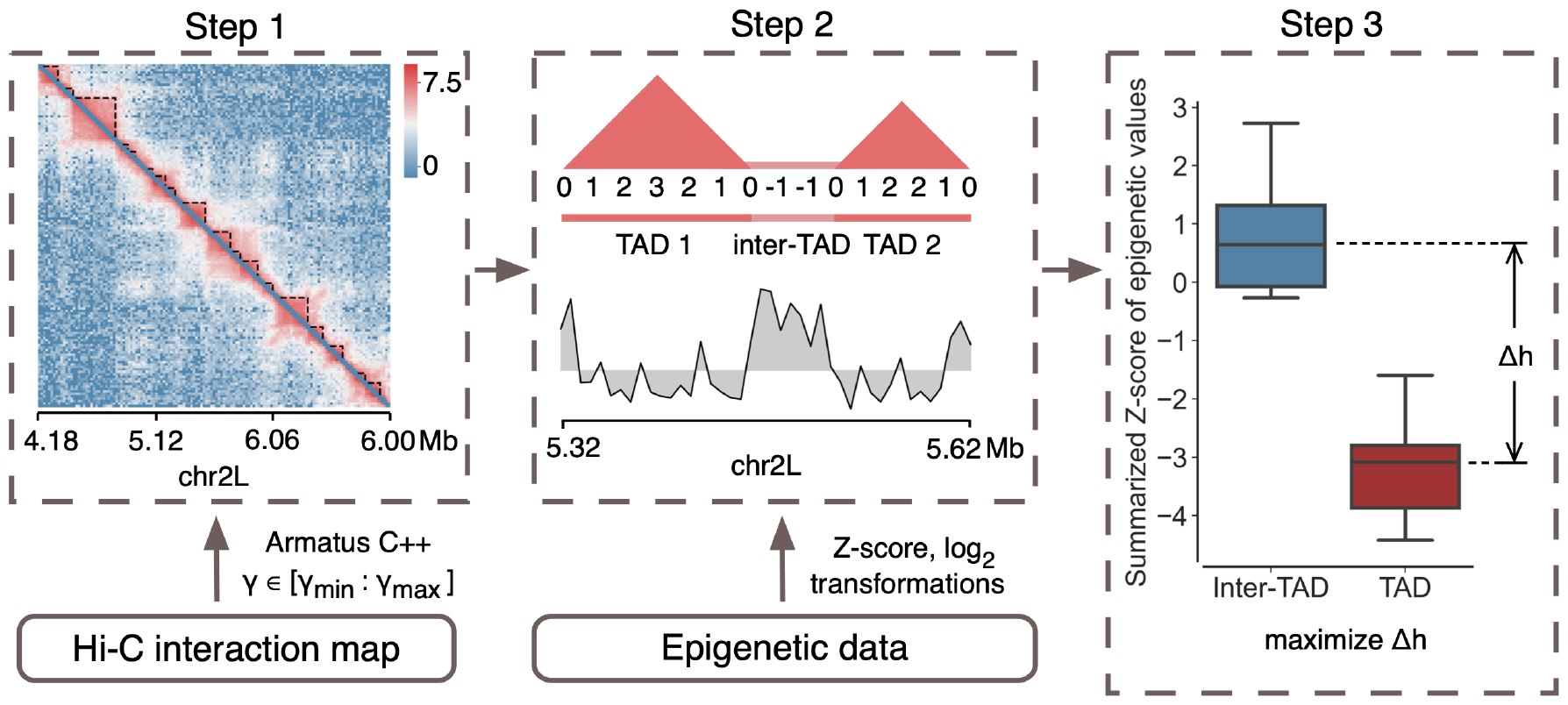
A general scheme of the `optimalTAD` algorithm. First, the algorithm takes a Hi-C matrix as an input and then identifies TAD regions for each value of the *γ* parameter from a predefined interval. Next, `optimalTAD` superimposes the obtained TAD sets onto binarized and scaled epigenetic profiles collected in ChIP-seq or similar experiments. Finally, it calculates the difference Δ*h* between epigenetic values within TADs and inter-TADs and optimizes the *γ* based on the maximal value of Δ*h*

### Annotation of TADs based on histone acetylation levels

To examine the ability of our algorithm to identify TAD sets under varying conditions, we tested `optimalTAD` using Hi-C and histone acetylation ChIP-seq data from *Ulianov et al*.. Specifically, we analyzed data from control S2 *Drosophila* cells and cells upon lamin Dm0 knock-down (Lam-KD). Our results showed that `optimalTAD` reported lower optimal *γ* values for control replicates (1.65 and 1.29) compared to Lam-KD replicates (2.15 and 1.73). Moreover, the acetylation amplitude curves for both Lam-KD replicates exhibited a specific shape, which differed noticeably from the curves for both control replicates (Fig. 2a). These distinct shapes provide a new opportunity for clustering analysis of Hi-C samples (Fig. 2b) because the acetylation amplitude curves reflect the epigenetic properties of TADs at different hierarchical levels.

**Fig. 2.**
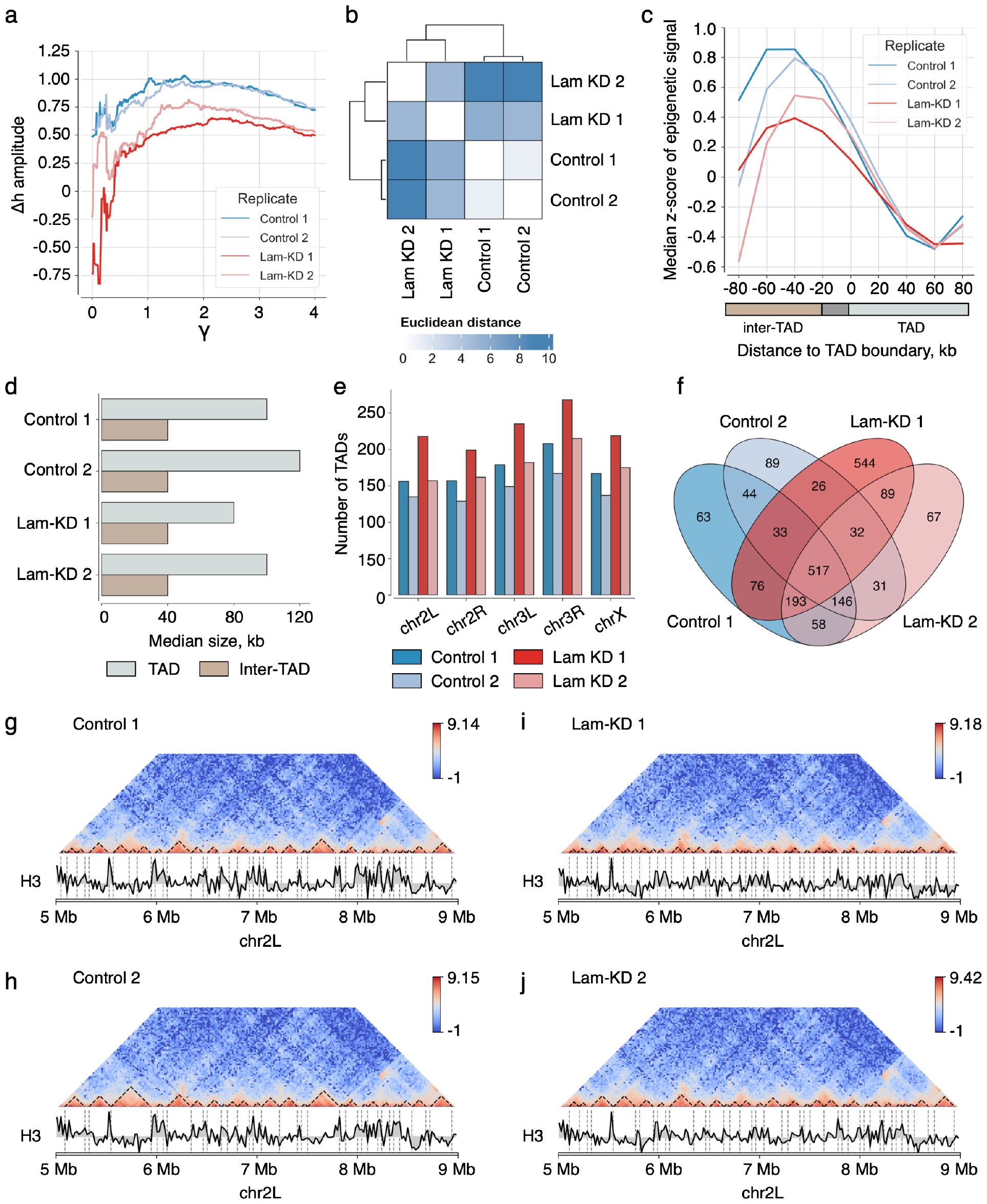
Algorithm validation using data from Ulianov et al., 2019. (a) Amplitude curves showing the dynamics of the difference Δ*h* between the average ChIP-seq signals within TADs and inter-TADs across various *γ* values. (b) Heatmap illustrating the hierarchical sample clusterization based on the Euclidean distance between the amplitude curves. (c) The amplitude of histone acetylation levels in the TAD set predicted by `optimalTAD` in the Ulianov dataset for *Drosophila*. (d) Comparison between TAD and inter-TAD median sizes obtained for experimental samples. (e) Distribution of TAD numbers predicted by `optimalTAD` on 2L, 2R, 3L, 3R, and X chromosomes. (f) Venn diagram showing the number of TAD boundaries shared between experimental replicates. (g-j) Hi-C maps for the genomic interval chr2L:5,000,000-9,000,000 showing the predicted optimal TADs (dashed lines) in control (panels g,h) and Lam-KD (panels i,j) replicates. Below the Hi-C maps, H3 ChIP-seq profiles for the same genomic interval are provided for comparison. In panels a,c,f, control replicates (Control-1, Control-2) are shown by shades of blue, while Lam-KD replicates (Lam-KD-1, Lam-KD-2) are represented by shades of red.

Next, we compared acetylation profiles near TAD boundaries obtained for the predicted optimal *γ* values in each replicate. As expected, we found a drastic difference in histone H3 acetylation levels between inter-TADs and TADs (Fig. 2c). In agreement with [35], inter-TAD histone acetylation enrichment decreased in Lam-KD. The median size of predicted TADs varied from 80 to 120 kb (80-100 kb for Lam-KD replicates and 100-120 kb for control replicates, consistent with their optimal *γ* values), while the median inter-TAD size remained constant across replicates and was equal to 40 kb (Fig. 2d). These observations agree with the *Drosophila* TAD properties reported in previous studies [38, 32].

Due to the substantial differences in the obtained Δ*h* amplitude curves and median TAD sizes between the Control and Lam-KD replicates, we further characterized a composition of the predicted TAD sets. While no TADs were detected on the 4th chromosome at different resolutions, other chromosomes comprised 135-242 topological domains, with the largest number of TADs observed in the Lam-KD-1 replicate on chromosome 3R (Fig. 2e). As expected, the predicted TADs were similar across experimental conditions, with 517 boundaries (out of 2008 detected in total) being shared among all four replicates (Fig. 2f). Other 193 TAD boundaries were predicted in all replicates except Control-2. However, certain TAD boundaries were condition-specific. In particular, we detected 89 boundaries that were control-specific and 44 boundaries that only appeared in Lam-KD samples. An example of TAD decomposition into two sub-TADs of smaller sizes in the Lam-KD samples is shown in Fig. 2g-j. Thus, the `optimalTAD` method can be successfully applied for the identification of differential TAD structures.

### Annotation of TADs based on architectural proteins

Further, we aimed to determine *Drosophila* architectural proteins and chromatin marks that were the most effective for predicting optimal TAD sets. We selected ChIP-Seq profiles of architectural proteins (CTCF, CP190, RAD21, BEAF-32, TFIIIC, Z4, L(3)mbt, Chromator, CBP, CapH2, RNAPII) together with chromatin mark profiles (H3K4me3, H3K9me2) and profiles of the Polycomb-group proteins (PC-RJ, PC-VP) from the Li dataset. Consistent with previous studies on predictors of TAD boundaries in *Drosophila* [32, 38, 3], our analysis revealed that RNAPII, BEAF32, and H3K4me3 were highly enriched near the predicted TAD boundaries (Fig. 3a,b) and exhibited maximal Δ*h* values (1.692, 1.532, and 1.527, respectively). On the contrary, CTCF was weakly enriched near TAD boundaries, indicating that it was far less suitable for the identification of TADs, in line with its limited role in TAD formation in flies [36, 17].

**Fig. 3.**
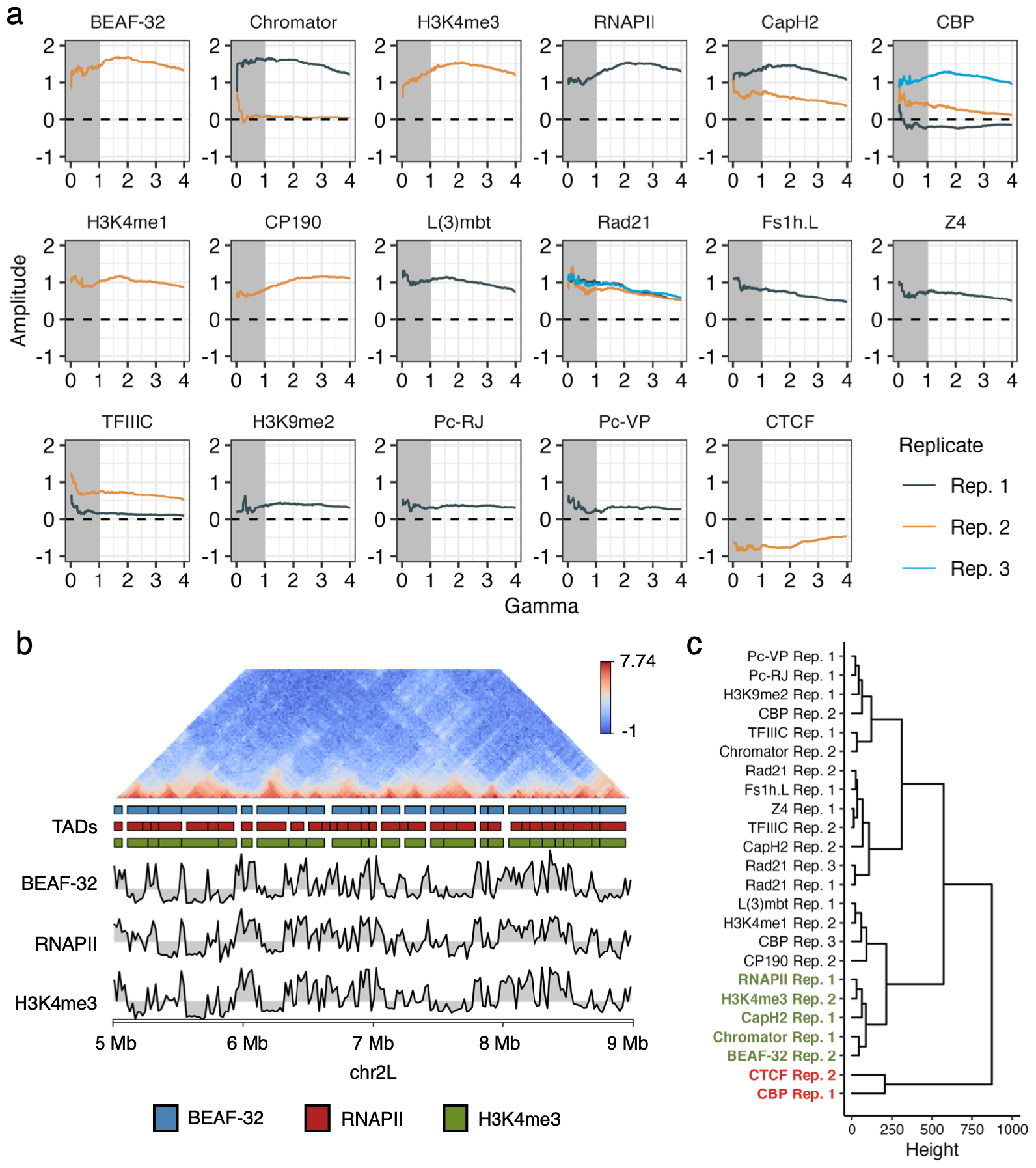
Algorithm validation using data from Li et al., 2015. (a) Amplitude curves showing changes in Δ*h* across different *γ* values for ChIP-seq replicates (Rep. 1-3, shown by shades of blue) of architectural proteins. (b) An example of the Hi-C map with TAD boundaries predicted in the genomic region chr2L:5,000,000-9,000,000 using BEAF-32 (blue rectangles), RNAPII (red rectangles), and H3K4me3 (green rectangles) profiles. (c) Hierarchical clusterization of the amplitude curves predicted using ChIP-seq data of different architectural proteins based on the Manhattan distance measure. Notable replicates producing maximal and minimal Δ*h* differences are highlighted by green and red rectangles, respectively.

Next, we hypothesized that the most suitable features would demonstrate similar changes in Δ*h* amplitude across different *γ* values. To test this hypothesis, we proceeded with the hierarchical clustering analysis of Δ*h* amplitude shapes obtained for the analyzed ChIP-seq replicates using the imputed Hi-C maps. Consistent with the results on the Δ*h* maximization (Table 1), the top five most suitable replicates (BEAF-32 Rep. 2, Chromator Rep. 1, H3K4me3 Rep. 2, RNAPII Rep. 1, CapH2 Rep. 1) formed a distinct cluster exhibiting consistent amplitude trends across the range of tested *γ* values (Fig. 3c).

**Table 1.** optimalTAD statistics obtained for the Li dataset using imputed and unimputed Hi-C data.

| Replicate | $\Delta h_{imp}$ | $\gamma_{imp}$ | $\Delta h_{unimp}$ | $\gamma_{unimp}$ |
| --- | --- | --- | --- | --- |
| BEAF-32 Rep. 2 | 1.69 | 1.7 | 1.36 | 2.66 |
| Chromator Rep. 1 | 1.68 | 1.16 | 1.23 | 2.89 |
| H3K4me3 Rep. 2 | 1.53 | 2.11 | 1.14 | 2.67 |
| RNAPII Rep. 1 | 1.53 | 2.1 | 1.22 | 2.65 |
| CapH2 Rep. 1 | 1.48 | 1.26 | 1.09 | 2.66 |
| CBP Rep. 3 | 1.29 | 1.81 | 0.89 | 3.54 |
| H3K4me1 Rep. 2 | 1.17 | 1.74 | 0.72 | 2.87 |
| CP190 Rep. 2 | 1.16 | 2.99 | 0.9 | 2.83 |
| L(3)mbt Rep. 1 | 1.14 | 1.57 | 0.78 | 2.49 |
| Rad21 Rep. 1 | 1.02 | 1.01 | 0.53 | 2.87 |
| Rad21 Rep. 3 | 1 | 1.34 | 0.6 | 2.87 |
| Rad21 Rep. 2 | 0.86 | 1.57 | 0.41 | 3.02 |
| Fs1h.L Rep. 1 | 0.84 | 1.14 | 0.48 | 2.52 |
| Z4 Rep. 1 | 0.79 | 1.16 | 0.39 | 2.53 |
| CapH2 Rep. 2 | 0.77 | 1.07 | 0.36 | 2.47 |
| TFIIIC Rep. 2 | 0.77 | 1.12 | 0.51 | 2.47 |
| CBP Rep. 2 | 0.45 | 1.01 | 0.13 | 2.53 |
| H3K9me2 Rep. 1 | 0.44 | 1.55 | 0.41 | 2.38 |
| Pc-RJ Rep. 1 | 0.38 | 1.87 | 0.24 | 3.56 |
| Pc-VP Rep. 1 | 0.34 | 1.64 | 0.19 | 3.57 |
| TFIIIC Rep. 1 | 0.16 | 1.42 | 0.04 | 3.8 |
| Chromator Rep. 2 | 0.11 | 1.07 | 0.01 | 4 |
| CBP Rep. 1 | -0.13 | 3.73 | -0.17 | 4 |
| CTCF Rep. 2 | -0.46 | 3.9 | -0.46 | 1.01 |

$\Delta h_{imp}$ and $\gamma_{imp}$ correspond to results with empty line imputation mode, $\Delta h_{noimp}$ and $\gamma_{noimp}$ represent results with unimputed Hi-C data

To evaluate the algorithm performance in the empty-line imputation mode, we tested it using data from [20]. Our results revealed that in runs with no imputation, the median value of the optimal *γ* was higher (2.82) compared to runs with imputed Hi-C (1.56) (Suppl. Fig. 1, Table 1). Conversely, the empty-line imputation mode resulted in a higher median size of predicted TADs, compared to those predicted for unimputed Hi-C maps. While handling missing bins in `optimalTAD` may help align the median TAD size with the typical TAD size in *Drosophila*, previously estimated as 100-150 kb [37], this approach also results in a substantial oscillation of the Δ*h* amplitude curve for *γ ∈* [0, 1] (Fig. 3a). In contrast, an optimal *γ >* 1 for the analysis with bin imputation produced a biologically reasonable and more stable median TAD size.

### Optimal TAD prediction in Toll^rm9/rm10^ mutant embryos

To illustrate the broad applicability of `optimalTAD`, we additionally tested it using the *Ing-Simmons et al*. dataset, which includes the Toll^rm9/rm10^ 16-kb Hi-C map and two H3K27ac ChIP-seq replicates. Our analysis resulted in two optimal TAD sets, both with a median size of 64 kb, and revealed that 92% of TAD boundaries were shared across the two domain sets (Fig. S2a,b). Additionally, the amplitude curves exhibited notable similarity (Suppl. Fig. S2c) with the Pearson’s R = 0.9942 and RMSD = 0.0018, highlighting the reproducibility of `optimalTAD` segmentations between biological replicates.

### Testing optimalTAD on individual chromosomes

The default method of *γ* optimization in `optimalTAD` calculates Δ*h* using information about TAD structure derived from all available chromosomes in *Drosophila*. This approach maximizes the number of ChIP-seq values used for *M* ^interTAD^ calculation (see Eq. 1 in Methods section) and enhances its robustness for low *γ* values. However, `optimalTAD` can be applied to analyze individual chromosomes. To demonstrate the algorithm performance in this context, we selected the ChIP-seq data for the top six features (BEAF-32, Chromator, H3K4me3, RNAPII, CapH2, CBP) exhibiting the highest Δ*h* values in the results presented above and analyzed 2L, 2R, 3L, 3R, and X chromosomes separately (Fig. 4). While the replicates demonstrated relatively low variability in maximal Δ*h* amplitude values across chromosomes (SD *∈* [0.06, 0.14]), they differed substantially in the values of optimal *γ* (SD *∈* [0.14, 1.29]). Only three tested replicates (BEAF-32 Rep. 2, CapH2 Rep. 2, and CBP Rep. 3) yielded a low-variable optimal *γ* (SD *<* 0.25). The optimal *γ* values for the chromosomes 3L, 3R, and 2L were the most consistent among replicates, with *γ ∈* [1.56, 1.62], *γ ∈* [1.8, 1.33], and *γ ∈* [1.85, 2.49], respectively (only the best replicate for each feature was considered). The highest variability was observed for *γ* values predicted for the chromosome 2R. These findings suggest that a combination of several proteins and chromatin marks might be potentially beneficial for the `optimalTAD` analysis.

**Fig. 4.**
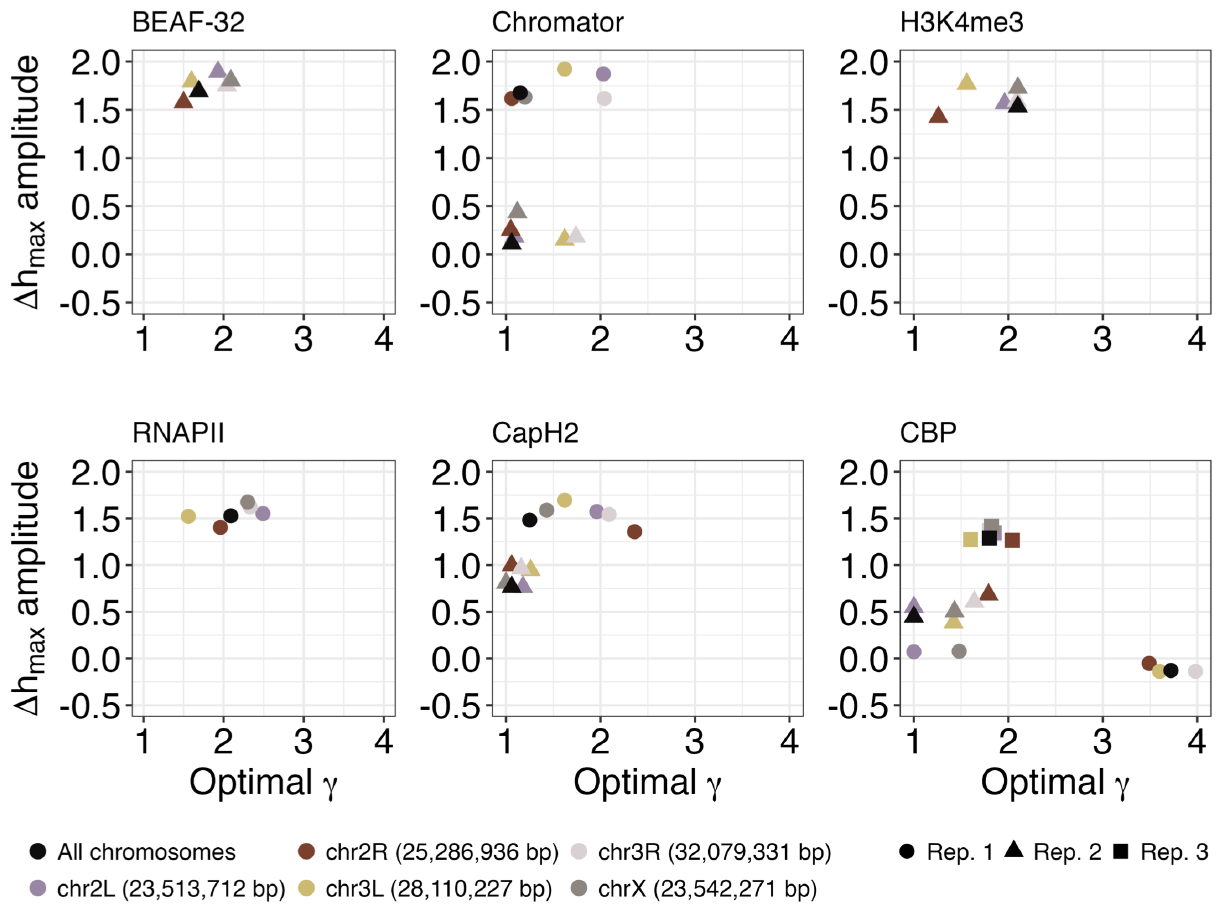
Distribution of optimal *γ* and Δ*h* values predicted for individual chromosomes (2L, 2R, 3L, 3R, and X chromosomes colored by purple, brown, yellow, pink, and grey colors, respectively), as well as for all chromosomes (colored by black). This analysis is based on the ChIP-seq profiles of the top six architectural proteins, which were the most suitable for the TAD identification. Results obtained for Rep. 1,2,3 are shown by circle, triangle, and rectangle shapes, respectively.

### Comparison with other TAD calling tools

Finally, we compared `optimalTAD` with several previously published TAD calling algorithms, such as `TopDom` [33], `DomainCaller` [39], `TADtree` [41], `OnTAD` [2], `ClusterTAD` [26], `Arrowhead` [29], `Caspian` [11], `HiCseg` [19]. All algorithms were tested using the 20-kb resolution Hi-C data from the *Li* study. Three tools (`TADtree`, `OnTAD`, and `Arrowhead`) produced multiple TAD sets in a hierarchy, making direct comparison with the TADs predicted by `optimalTAD` not feasible. Two other tools (`DomainCaller (TADlib)` and `Arrowhead`) yielded non-hierarchical TAD sets but without inter-TAD regions characteristic for the `Drosophila` genome. Therefore, we proceeded with the three remaining algorithms (`TopDom`, `ClusterTAD`, and `Caspian`) using their default parameters.

To evaluate the performance of each TAD calling method, we calculated the Δ*h* amplitude value using the BEAF-32 ChIP-seq profile. Among these algorithms, `optimalTAD` showed the highest value of Δ*h*, outperforming other algorithms by more than 1.7-fold (Figure 5a). Consistently, the acetylation curve predicted using `optimalTAD` showed the most dramatic difference in acetylation levels between inter-TAD and TAD regions, compared to other algorithms (Fig. 5b-e).

**Fig. 5.**
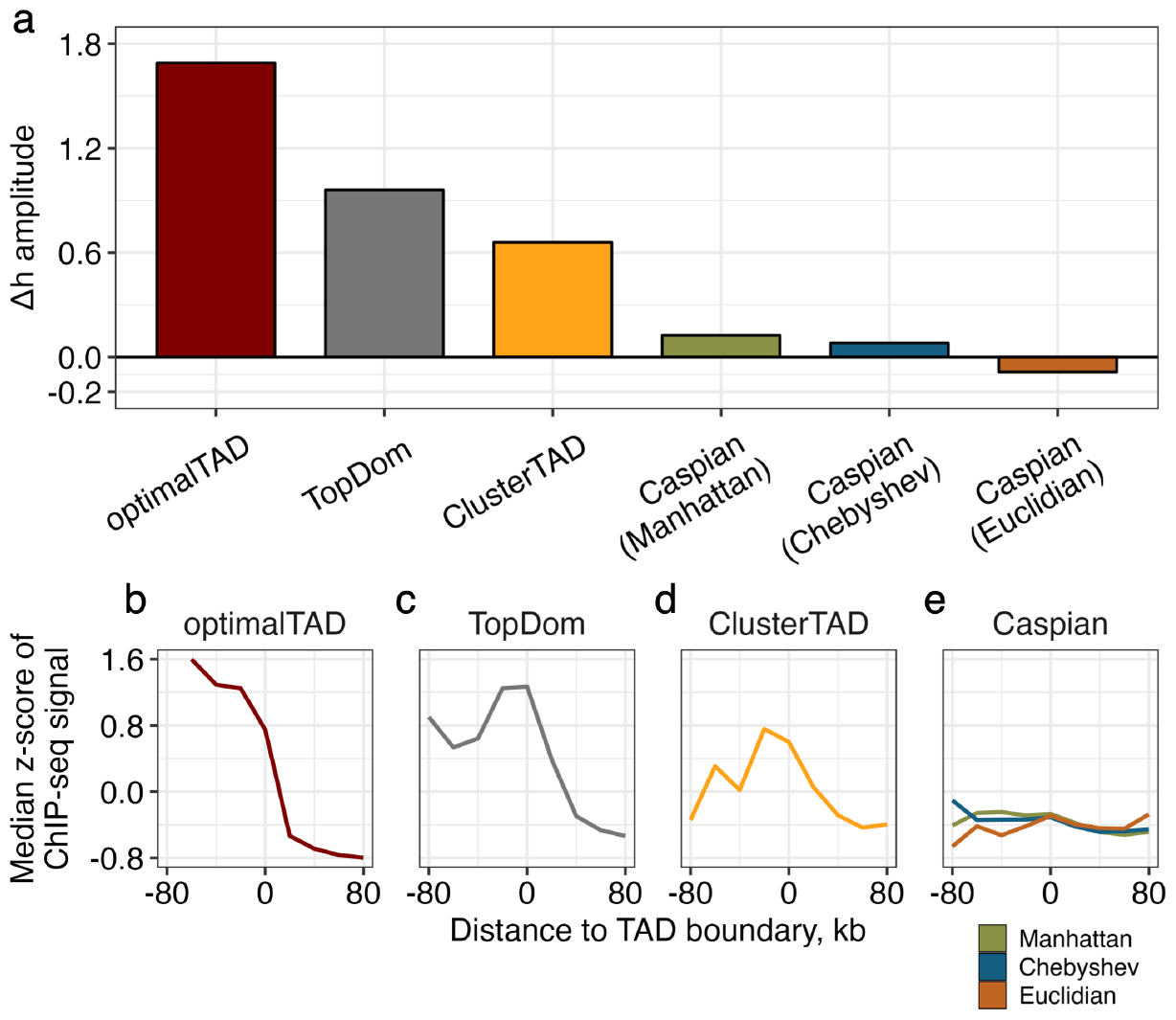
Comparison of `optimalTAD` with other TAD calling tools. (a) Barplot showing Δ*h* amplitude values calculated using `optimalTAD, TopDom, ClusterTAD`, and `Caspian` methods (from left to right) for the Hi-C map from Li dataset. An analysis with the Caspian tool was done in three possible modes: “Manhattan”, “Chebyshev”, and “Euclidean”. (b-e) Comparison of histone acetylation curves corresponding to TAD calling tools selected for the analysis.

### Algorithm validation using mammalian data for neurons and glial cells

*Drosophila* and mammalian TADs are essentially different in several key characteristics [7, 23, 38]. The most prominent dissimilarity lies in the mechanism of TAD formation. In fruit flies, TADs exhibit a compartmental nature, while in mammals, there are both loop-extrusion based domains and compartmental ones [21, 25, 31]. Additionally, the nature of TADs is influenced by the diversity of architectural proteins occupying domain boundaries, with BEAF32 and CTCF being predominant at *Drosophila* and mammalian TAD boundaries, respectively [34, 13]. Furthermore, while mammalian TADs are continuously organized, *Drosophila* TADs are separated by inter-TADs.

Yet, despite all these differences in the fundamental principles of TAD organization, the family of TAD annotation algorithms suitable for mammals and relying heavily on the strength of insulation between adjacent loci also have one parameter that regulates the size and total number of TADs. Therefore, similar to *Drosophila*, `optimalTAD` is applicable for TAD annotation in mammals, although it must utilize specific TAD callers capable of capturing chromatin domains that are distinct from *Drosophila* ones. In addition, Δ*h* calculations should be based on species-specific architectural proteins relevant to the TAD organization in a particular species.

Aiming to validate the applicability of `optimalTAD` to mammalian TADs, we analyzed publicly available Hi-C data for glial and neuronal chromatin structure in the mouse cortex [4]. We also used ChIP-seq and meDIP-seq tracks for several key histone marks and DNA methylation generated for the same types of cell populations that were obtained from the anterior cingulate cortex [12]. Additionally, we tested the tool on the CTCF ChIP-seq track available for cortical neurons only [16].

The `optimalTAD` algorithm was applied to mouse data, resulting in a set of boundaries for glial and neuronal Hi-C maps at 80- and 60-kb Insulation Score (IS) windows (see Methods), respectively, which were automatically selected as optimal by the algorithm. The boundaries demonstrated classic insulating properties (Fig. 6b-c, bottom) with a striking spike of CTCF (Fig. 6b, top), consistent with its well-known architectural role [7, 18]. The number and size of annotated TADs fell within the ranges reported in previous studies of chromatin organization in mouse neurons [16, 22].

**Fig. 6.**
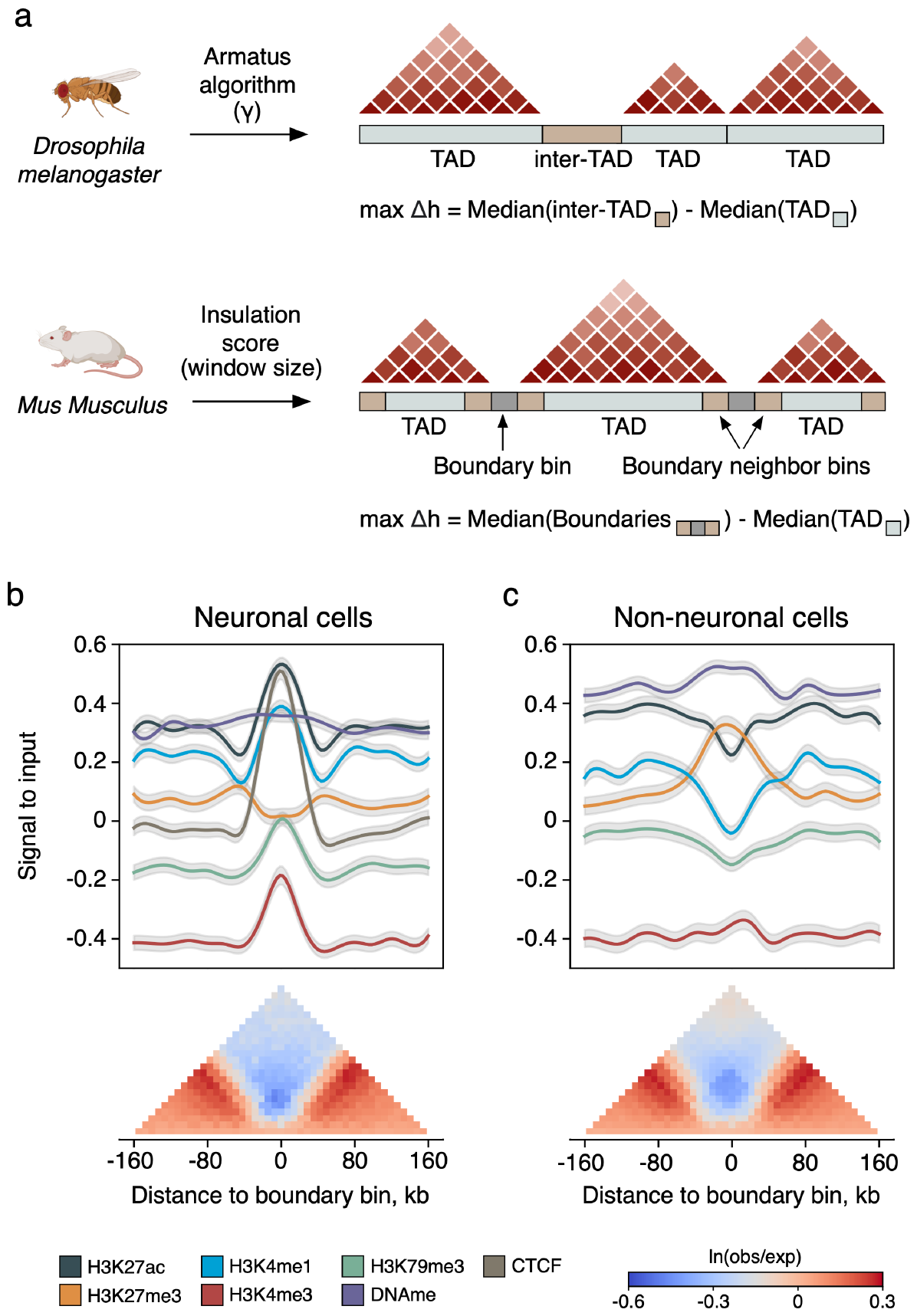
`optimalTAD` successfully predicts an optimized set of TAD boundaries in the mouse cortex. (a) A generalized scheme summarizing the distinct optimization strategies implemented in `optimalTAD` for *Drosophila* and mammalian data. The *Drosophila* genome contains both TAD and inter-TAD regions, while the mammalian genome is partitioned into continuous topological domains. Therefore, for mammalian data, `optimalTAD` considers boundary bins and adjacent bins instead of inter-TADs. Then, it optimizes the window size in the Insulation Score (IS) algorithm (see Methods). (b-c) Top: tracks of histone marks and DNA methylation around TAD borders detected with `optimalTAD` on data for neuronal (b) and glial (c) cell populations. Red, blue, black, green, orange and purple curves correspond to H3K4me3, H3K4me3, H3K27ac, H3K79me3, H3K27me3 histone modifications and DNA methylation profile, respectively. The gray curve represents CTCF profile available only for neurons. Lines represent median values of log_2_-normalized z-score of ChIP-seq signals divided by inputs (see Methods). Bottom: average heatmaps generated around TAD boundaries annotated by `optimalTAD` in neuronal and glial Hi-C maps.

Interestingly, the distribution of histone marks around TAD boundaries revealed fundamental differences between neurons and glial cells. In particular, histone marks representing active genes and enhancers were abundant at the TAD boundaries detected for neuronal Hi-C data (Fig. 6b, top), while a similar analysis conducted for glial chromatin maps revealed an increase in repressive mark occupancy, consistent with current understanding of chromatin organization in these cell types [28]. Specifically, we observed a pronounced enrichment of Polycomb-associated H3K27me3 marks at the TAD boundaries in glial cells, which was also supported by DNA methylation curves (Fig. 6c, top). Moreover, ChIP-seq signals of active enhancers (H3K27ac, H3K4me1) were strongly depleted at the TAD boundaries in glial cells (Fig. 6c, top).

## Discussion

Despite many approaches developed, the computational identification of TADs remains challenging due to their nested structure, leading to ambiguity in the TAD annotation. Identified TADs are commonly evaluated by visual inspection of Hi-C maps, which impairs the reproducibility of chromatin architecture studies. The definition of TADs is highly dependent on the scale and resolution at which they are searched. On the same Hi-C map, it is possible to annotate megabase- and kilobase-scale TAD sets, both of which appear to be visually valid.

In this study, we address this issue by developing the `optimalTAD` algorithm that combines Hi-C and epigenetics (such as ChIP-seq) data for robust TAD annotation. TAD boundaries are typically enriched with specific chromatin marks, which creates a pronounced gradient of architectural proteins, histone acetylation and other chromatin marks at TAD boundaries. `optimalTAD` identifies multiple TAD sets at different scales in the Hi-C map and then selects the best TAD segmentation with the maximum enrichment of histone marks or architectural proteins at TAD boundaries. In addition, `optimalTAD` identifies epigenetic marks that are best suited for TAD prediction.

Therefore, `optimalTAD` incorporates functional information about histone marks and architectural proteins to annotate TADs as both structural and functional units of chromatin. We compare `optimalTAD` with existing approaches that are limited to using only structural information about contact frequencies and predict boundaries based on a predefined TAD size. `optimalTAD` surpasses these approaches.

Our analysis of multiple *Drosophila* and mouse datasets suggests that `optimalTAD` is a universal tool suitable for both flies and mammals. Despite very distinct TAD organization principles in these groups of organisms, `optimalTAD` produces correct TAD annotations in each of the many datasets we considered, including drastically different cell types with highly specialized profiles of chromatin marks enriched at TAD boundaries. The quality of the resulting TAD annotations allows drawing valid conclusions about TAD structure, functions, and properties that are specific for a particular cell type and organism and align well with previous studies.

## Conclusions

In conclusion, `optimalTAD` provides an optimized identification of topological domains and can be successfully applied to Hi-C data obtained from various sources. While other TAD-calling algorithms are limited to using only structural information on contact frequencies and predict boundaries based on a predefined TAD size, `optimalTAD`, in addition, incorporates functional information about chromatin marks and architectural proteins to annotate TADs as both structural and functional units of chromatin. Most importantly, `optimalTAD` solves the problem of arbitrary TAD identification. Due to the high cost of Hi-C experiments and the complexity of Hi-C analysis, approaches improving the reproducibility of Hi-C analysis are essential for researchers performing chromatin conformation studies.

## Methods

### The optimalTAD algorithm

- **Input data**. Hi-C contact maps should be normalized using the iterative correction method [14]. ChIP-seq profiles should be normalized using both the total number of reads and corresponding inputs, and further log_2_ and Z-score transformed. `optimalTAD` binarizes ChIP-seq coverage into intervals of the same length as the resolution of the Hi-C contact matrix.
- **Empty line imputation**. A typical Hi-C matrix includes rows and columns consisting entirely of missing values. Such empty lines are often found in poorly mapped genome regions and areas enriched with repeated DNA sequences. These empty lines hinder the accurate TAD identification. To address this issue, `optimalTAD` utilizes linear regression to impute empty bins within such lines based on neighboring bins, thus facilitating improved TAD annotation.
- **TAD calling**. The next step of TAD set identification can be carried out in two possible modes, depending on whether the input Hi-C data is from *Drosophila* or mammalian species. For *Drosophila* Hi-C data, TAD calling is performed using the `Armatus` tool for each resolution parameter *γ ∈* [*γ*_*min*_, *γ*_*max*_] with a fixed step *s*. For mammalian Hi-C data, TAD boundaries are identified using the Insulation Score method implemented in the `cooltools` package [27] with a predefined set of insulation window sizes *ω ∈* [*ω*_*min*_, *ω*_*max*_].
- **TAD indexing**. Once all TAD sets for the input Hi-C map are determined, `optimalTAD` performs inter- and intra-domain bin indexing. Each bin index is calculated as the distance from the bin to the nearest TAD boundary (Fig. 1a, step 2). Bins corresponding to domain boundaries are indexed as 0, while inter-TAD bin indices are multiplied by *−*1. Next, each bin index is matched with the corresponding value from the binarized epigenetic profile (Fig. 1a, step 3). Finally, for each TAD set, `optimalTAD` calculates the medians *M* ^TAD^ and *M* ^interTAD^ of ChIP-seq values belonging to TAD and inter-TAD regions, respectively.
- **Optimization**. We next optimize the TAD calling parameter *γ* by maximizing the difference Δ*h* between the median ChIP-seq values in inter-TADs and TADs:

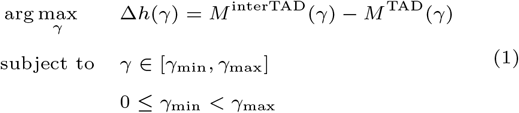

where *γ* is the resolution parameter in `Armatus`, while *M* ^interTAD^ and *M* ^TAD^ are the medians of ChIP-seq values in inter-TAD and TAD bins, respectively.

For mammalian data, *M* ^interTAD^ in Eq. 1 represents the median of epigenetic signals from TAD boundary bins (S_*b*_) and their adjacent bins (S_*b−*1_ and S_*b*+1_). The optimized variable *γ* is replaced with the insulation window size *ω*. Thus, the best window size *ω* is determined by maximizing the difference in median epigenetic signals between expanded boundary bins (S_*i*_, *i ∈* [*b −* 1, *b* + 1]) and TAD bins (S_*i*_, *i ∈/* [*b −* 1, *b* + 1]) (Fig. 6a).

### Implementation

To accelerate the TAD calling step, we developed a parallel implementation of the `Armatus` algorithm that supports *>* 1 computational core. Parallel `Armatus` decomposes multiscale domain segmentation into *N* threads, where *N* is equal to the number of cores available on the machine. Therefore, it achieves maximal computational efficiency when using 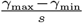 cores.

### Algorithm validation

Three publicly available datasets [35, 20, 15] were used for the algorithm validation in *Drosophila*. We used 20-kb Hi-C maps from the Ulianov and Li datasets, along with 16-kb Hi-C maps from the Ing-Simmons dataset. To standardize the Hi-C interaction frequency values, we scaled them from *x*_min_ = 1 to *x*_max_ = 1024 and then log_2_ transformed. The log transformation was implemented to bring the distribution of values closer to the normal distribution. Processed ChIP-seq profiles were also log_2_ transformed and standardized using Z-score. All `optimalTAD` validation runs were done with the step size parameter *s* = 0.01 using four cores on MacOS Catalina (Intel Core i5).

To demonstrate the applicability of `optimalTAD` for mammalian genome analysis, we utilized publicly available Hi-C [4] and epigenetic (ChIP-seq, meDIP-seq) [12, 16] datasets of mouse cortical neurons (NeuN+) and non-neuronal (NeuN-) cells. 20-kb resolution Hi-C maps were chosen for the analysis. The window sizes in the IS algorithm varied from 40 to 200 kb, with a step size of 20 kb. Chromosomes M, X, Y and detected TADs larger than 2 Mb were discarded by the algorithm. Boundaries corresponding to bins with low coverage (*”is bad bin” = True*) were also excluded from the prediction.

### Data preprocessing

#### Ulianov et al., 2019

Processed Hi-C and ChIP-seq data of two controls and two lamin Dm0 knock-down *Drosophila* S2 cell line replicates were obtained from the GEO repository (GSE110082 accession number). H3 ChIP-seq experiments were normalized by their respective inputs and then binned into 20-kb intervals using the following formula:

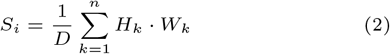

where *S*_*i*_ is a signal calculated for *i*-th bin, *D* is the bin size (20 kb), *H*_*k*_ is the peak height, *W*_*k*_ is the peak width, and *n* is the number of peaks.

#### Li et al., 2015

This dataset features both Hi-C interaction maps and ChIP-seq profiles of various architectural proteins obtained for the *Drosophila* KC167 cell line under heat shock and normal conditions. Raw Hi-C reads of samples without treatment were downloaded from GEO (GSE62904) and then processed using `Distiller` [1]. Similar to the first dataset, downloaded ChIP-seq replicates were normalized by inputs and partitioned into 20-kb regions according to the Eq. 1.

#### Ing-Simmons et al., 2021

Raw ChIP-seq reads for H3K27ac of Tollrm9/rm10 replicates were downloaded from ArrayExpress (accession number E-MTAB-9303) and then processed using the `nf-core/chipseq` pipeline (version 1.2.1). Alignments with low mapping quality (MAPQ *<* 30) were filtered out. The ChIP-seq signal was calculated within each 0.1-kb bin and normalized by the total number of reads in the library. The Hi-C interaction maps of Tollrm9/rm10 mutants were also obtained from ArrayExpress (E-MTAB-9306).

#### Hi-C and ChIP-seq data for mouse neuronal and glial cortical cells

Processed Hi-C data were downloaded from GSE168524 in `hic` format. They were converted to `cool` files with `hic2cool` tool, pooled among male and female samples at 20-kb resolution, and downsampled to the same total number of contacts for neurons and glia using `cooltools` [27]. To calculate average heatmaps around TAD boundaries, the obtained Hi-C maps were normalized using observed-over-expected approach and log-transformed. ChIP-seq tracks for histone modifications and the meDIP-seq track for DNA methylation were downloaded from GSE74964 in `bigwig` format. In this study, files for epigenetic modifications labeled as ‘naive’ have been used, considered as preceding any intervention. The tracks were merged among replicates. Then, each track was binned at 20-kb resolution and the signals obtained for histone modifications were divided by their corresponding inputs. For the CTCF track, the procedure was the same except for the initial stage: the ChIP-seq data for wild-type neurons were downloaded as raw reads from GSE99363 and processed with the `nf-core/chipseq` pipeline [9] using mm10 genome assembly.

### Description of TAD calling tools used for comparative analysis

- **TopDom** [33]. To perform the `TopDom` analysis, we used the R package available at https://github.com/jasminezhoulab/TopDom (version 0.0.2). The program was run with “window.size” = 10.
- **ClusterTAD** [26]. The Java version of the algorithm (version 1.0) was downloaded from https://github.com/BDM-Lab/ClusterTAD. TAD sets with the best quality score were considered as the optimal ones and then used to evaluate Δ*h*.
- **Caspian** [11]. For the TAD identification with `Caspian` tool, we used all three available metrics (”Euclidian”, “Manhattan”, and “Chebyshev”) implemented in this algorithm for the pairwise bin distance calculation. The algorithm was downloaded from https://gitee.com/ghaiyan/caspian and the default parameters were specified.

## Funding

This work was supported by the Russian Science Foundation (grant number 21-74-10102); and the Russian Foundation for Basic Research (project number 21-34-70051).

## Competing interests

No competing interest is declared.

## Author contributions statement

D.N.S. developed `optimalTAD` and analyzed data, A.D.K. performed validation, D.T. and E.E.K. supervised the analysis, M.S.G. and E.E.K. conceived the study. All authors contributed to writing the manuscript text.

**Dmitrii N. Smirnov** is a junior research scientist at the Skolkovo Institute of Science and Technology, Russia. He is also affiliated with the Ben-Gurion University of the Negev, Israel. His research interests include bioinformatics, the development of new computational methods for studying chromatin organization and the biology of aging.

**Anna D. Kononkova** is a junior research scientist at the Skolkovo Institute of Science and Technology, Russia. She was also affiliated with the Institute for Information Transmission Problems of the Russian Academy of Sciences, Moscow. She specializes in chromatin structural biology focusing on the computational analysis of Hi-C data.

**Debra Toiber** is an associate professor at the Ben-Gurion University of the Negev, Israel. Her research interests encompass neurobiology, sirtuin biology and molecular mechanisms behind ageing and age-related diseases.

**Mikhail S. Gelfand** is a professor at the Skolkovo Institute of Science and Technology, Russia. He is also affiliated with the Institute for Information Transmission Problems of the Russian Academy of Sciences, Russia. His research interests are in the areas of molecular evolution, comparative genomics and systems biology.

**Ekaterina E. Khrameeva** is an assistant professor at the Skolkovo Institute of Science and Technology, Russia. She specializes in developing computational methods and studying genome folding principles in a wide range of organisms, from unicellular ones to humans.

## Notes

### Competing Interest Statement

The authors have declared no competing interest.

### Summary of Updates

The manuscript was updated with 'Algorithm validation using mammalian data for neurons and glial cells' and 'Discussion' sections; old supplementary figures are included in the main text now.

