## Supplementary figures for "optimalTAD: annotation of topologically associating domains based on chromatin marks enrichment"

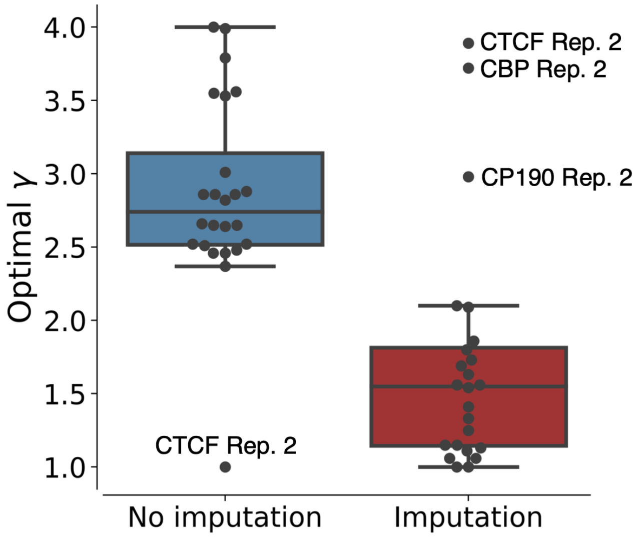


**Fig. 1.** Empty line imputation mode in the `optimalTAD` algorithm. Boxplots illustrate the distribution of the predicted $\gamma$ values in `optimalTAD` runs with empty bin imputation (red box) compared to runs with no empty bin imputation (blue box).

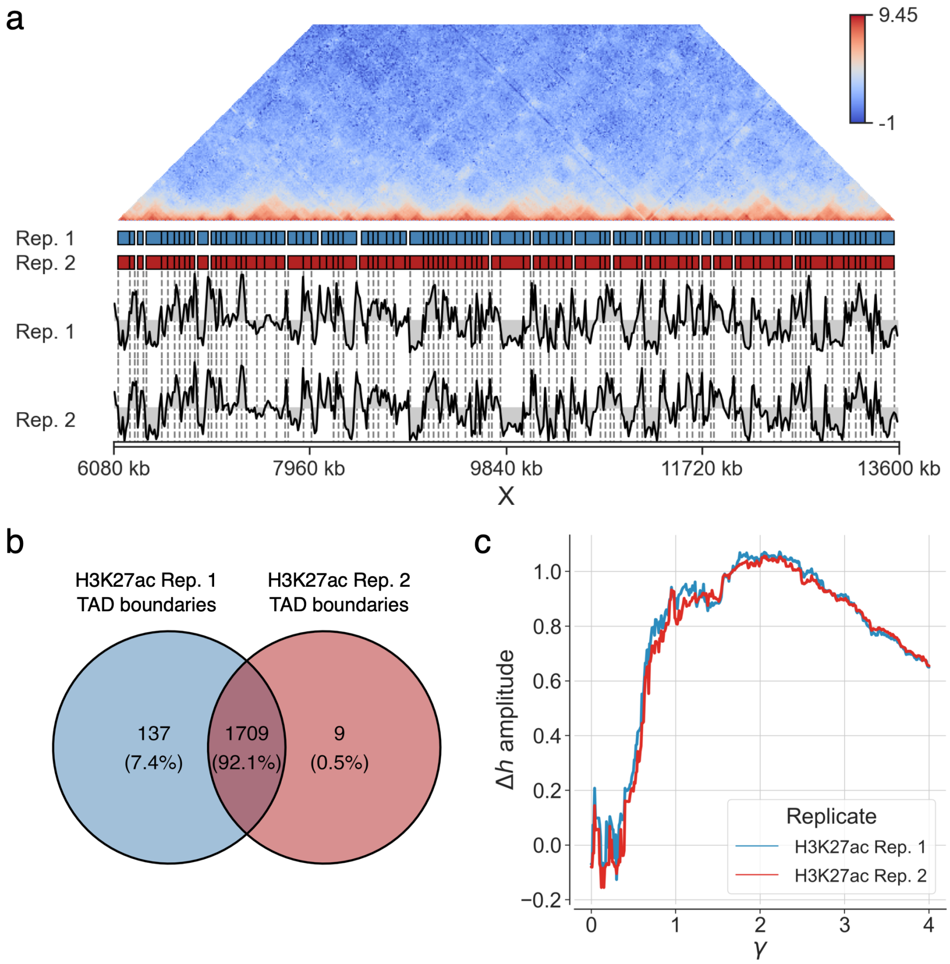


**Fig. 2.** Algorithm validation using Toll$^{rm9/rm10}$ mutant embryos data. (a) Amplitude curves showing changes in $\Delta h$ across $\gamma$ values for Rep. 1 and Rep. 2 of H3K27ac ChIP-seq profiles. (b) Venn diagram illustrating a number of shared boundaries between TADs predicted in two H3K27ac replicates (blue circle corresponds to Rep. 1 and the red circle shows Rep. 2 domains). (c) Hi-C map with TAD boundaries predicted for H3K27ac Rep. 1 (colored by blue) and H3K27ac Rep. 2 (colored by red) within the chrX:6,080,000-9,000,000 genomic interval.
